# A bioinformatics tool for identifying intratumoral microbes from the ORIEN dataset

**DOI:** 10.1101/2023.05.24.541982

**Authors:** Cankun Wang, Anjun Ma, Megan E. McNutt, Rebecca Hoyd, Caroline E. Wheeler, Lary A. Robinson, Carlos H.F. Chan, Yousef Zakharia, Rebecca D. Dodd, Cornelia M. Ulrich, Sheetal Hardikar, Michelle L. Churchman, Ahmad A. Tarhini, Eric A. Singer, Alexandra P. Ikeguchi, Martin D. McCarter, Nicholas Denko, Gabriel Tinoco, Marium Husain, Ning Jin, Afaf E.G. Osman, Islam Eljilany, Aik Choon Tan, Samuel S. Coleman, Louis Denko, Gregory Riedlinger, Bryan P. Schneider, Daniel Spakowicz, Qin Ma

## Abstract

Evidence supports significant interactions among microbes, immune cells, and tumor cells in at least 10–20% of human cancers, emphasizing the importance of further investigating these complex relationships. However, the implications and significance of tumor-related microbes remain largely unknown. Studies have demonstrated the critical roles of host microbes in cancer prevention and treatment responses. Understanding interactions between host microbes and cancer can drive cancer diagnosis and microbial therapeutics (bugs as drugs). Computational identification of cancer-specific microbes and their associations is still challenging due to the high dimensionality and high sparsity of intratumoral microbiome data, which requires large datasets containing sufficient event observations to identify relationships, and the interactions within microbial communities, the heterogeneity in microbial composition, and other confounding effects that can lead to spurious associations. To solve these issues, we present a bioinformatics tool, MEGA, to identify the microbes most strongly associated with 12 cancer types. We demonstrate its utility on a dataset from a consortium of 9 cancer centers in the Oncology Research Information Exchange Network (ORIEN). This package has 3 unique features: species-sample relations are represented in a heterogeneous graph and learned by a graph attention network; it incorporates metabolic and phylogenetic information to reflect intricate relationships within microbial communities; and it provides multiple functionalities for association interpretations and visualizations. We analyzed 2704 tumor RNA-seq samples and MEGA interpreted the tissue-resident microbial signatures of each of 12 cancer types. MEGA can effectively identify cancer-associated microbial signatures and refine their interactions with tumors.

**SIGNIFICANCE:** Studying the tumor microbiome in high-throughput sequencing data is challenging because of the extremely sparse data matrices, heterogeneity, and high likelihood of contamination. We present a new deep-learning tool, microbial graph attention (MEGA), to refine the organisms that interact with tumors.

## INTRODUCTION

The study of microbial communities and their impact on human health has gained increasing attention over the past decade (1). The role of intratumoral microbes in the tumor microenvironment has become an increasingly important area in studying the development and progression of cancer (2). The intratumoral microbiome affects outcomes in several cancers, including *Fusobacterium nucleatum* in the development of colon cancer and *Helicobacter pylori* in stomach cancer. To explore the relationship between the microbiome and cancer, large-scale genomic datasets such as The Cancer Genome Atlas (TCGA) have been utilized. However, limited attention has been given to the cancer-specific gene-microbe relationships. In this context, the Oncology Research Information Exchange Network (ORIEN) provides a real-world dataset consisting of clinical, genomic, and transcriptomic data collected under an institutional review board (IRB)-approved common protocol known as Total Cancer Care. It represents a valuable resource for identifying intratumoral microbes from various cancer types (3). Advances in sequencing technologies have provided large-scale human tissue sequencing data, which enables the characterization of the tissue-resident metagenome. However, exploring the links between the intratumoral microbiome and cancer tissues is ongoing due to the difficulties in obtaining clinical biopsies specifically dedicated to microbial profiling.

Here, we present Microbial Heterogeneous Graph Attention (MEGA), a deep learning-based Python package for identifying cancer-associated intratumoral microbes. The model is trained on ORIEN intratumoral microbial RNA sequencing (RNA-seq) data to identify microbial communities associated with each of the 12 human cancer types. The core framework is a heterogeneous graph transformer (HGT) (4) that can learn the importance and contribution of species to cancer samples. We have shown the superior performance of HGT in characterizing cell-gene relations from single-cell multi-omics datasets (5) and identifying sample-species relations (6) from The Cancer Microbiome Atlas (TCMA) data (7). To demonstrate the effectiveness and credibility of MEGA on the more complicated ORIEN data, we focus on 2 widely studied cancer types: colon adenocarcinoma (COAD) and thyroid carcinoma (THCA). By leveraging metabolic and phylogenetic relationships, MEGA was able to capture the association of low attention score microbes, suggesting the importance of integrating multiple types of data in identifying cancer-associated microbes. We believe that MEGA offers a comprehensive and nuanced approach to identifying cancer-associated intratumoral microbes in the ORIEN dataset, which could ultimately serve as potential targets for further study and therapy development.

## METHODS

### Study Design

Established in 2014, the Oncology Research Information Exchange Network (ORIEN) is an alliance of 18 US cancer centers. All ORIEN alliance members utilize a standard IRB-approved protocol: Total Cancer Care® (TCC). As part of the TCC, participants agree to have their clinical data followed over time, to undergo germline and tumor sequencing, and to be contacted in the future by their provider if an appropriate clinical trial or other study becomes available (8). TCC is a prospective cohort study where a subset of patients elect to be enrolled in the ORIEN Avatar program, which provides research use only (RUO)-grade whole-exome tumor sequencing, RNA-seq, germline sequencing, and collection of deep longitudinal clinical data with lifetime follow-up. Nationally, over 325,000 participants have enrolled in TCC. M2GEN, the commercial and operational partner of ORIEN, harmonizes all abstracted clinical data elements and molecular sequencing files into a standardized, structured format to enable the aggregation of de-identified data for sharing across the network. Data access was approved by the IRB in an Honest Broker protocol (2015H0185) and Total Cancer Care protocol (2013H0199) in coordination with M2GEN and participating ORIEN members.

### Sequencing Methods

ORIEN Avatar specimens undergo nucleic acid extraction and sequencing at HudsonAlpha (Huntsville, AL) or Fulgent Genetics (Temple City, CA). For frozen and OCT tissue DNA extraction, Qiagen QIASymphony DNA purification is performed, generating a 213 bp average insert size. For frozen and OCT tissue RNA extraction, Qiagen RNAeasy plus mini kit is performed, generating 216 bp average insert size. For formalin-fixed paraffin-embedded (FFPE) tissue, a Covaris Ultrasonication FFPE DNA/RNA kit is utilized to extract DNA and RNA, generating a 165 bp average insert size. RNA-seq is performed using the Illumina TruSeq RNA Exome with single library hybridization, cDNA synthesis, library preparation, and sequencing (100 bp paired reads at Hudson Alpha, 150 bp paired reads at Fulgent) to a coverage of 100M total reads/50M paired reads.

### Microbe Abundance and Diversity

RNA-seq reads are used to calculate microbe abundances using the {exotic} pipeline, as described previously (3). Briefly, reads are aligned first to the human reference genome, and then unaligned reads are mapped to a database of bacteria, fungi, archaea, viruses, and eukaryotic parasites. The observed microbes then proceed through a series of filtering steps to carefully and conservatively remove contaminants before batch correction and normalization. Diversity measures were estimated by calculating the Shannon and Simpson indices, as well as Chao1, ACE, and inverse Simpson using the R package vegan.

The input dataset for MEGA includes the microbiome matrix and the sample metadata of the cancer types. The raw counts of the ORIEN microbiome matrix consist of 2603 species in 2891 samples. The sample metadata is a two-column matrix that describes the label of the total of 12 cancer types at each sample. The NJS16 metabolic database (9) is a literature-curated interspecies network of the human gut microbiota, composed of approximately 570 microbial species and 3 human cell types metabolically interacting through more than 4400 small-molecule transport and macromolecule degradation events. We utilized the R package *taxizedb* to access the National Center for Biotechnology Information (NCBI) taxonomy database (10). It was integrated to prepare for the taxonomy ID to taxonomy name conversion and to extract additional phylogenetic relationships from the ORIEN data **(see Figure 1 – Data Sources)**.

**Figure 1.**
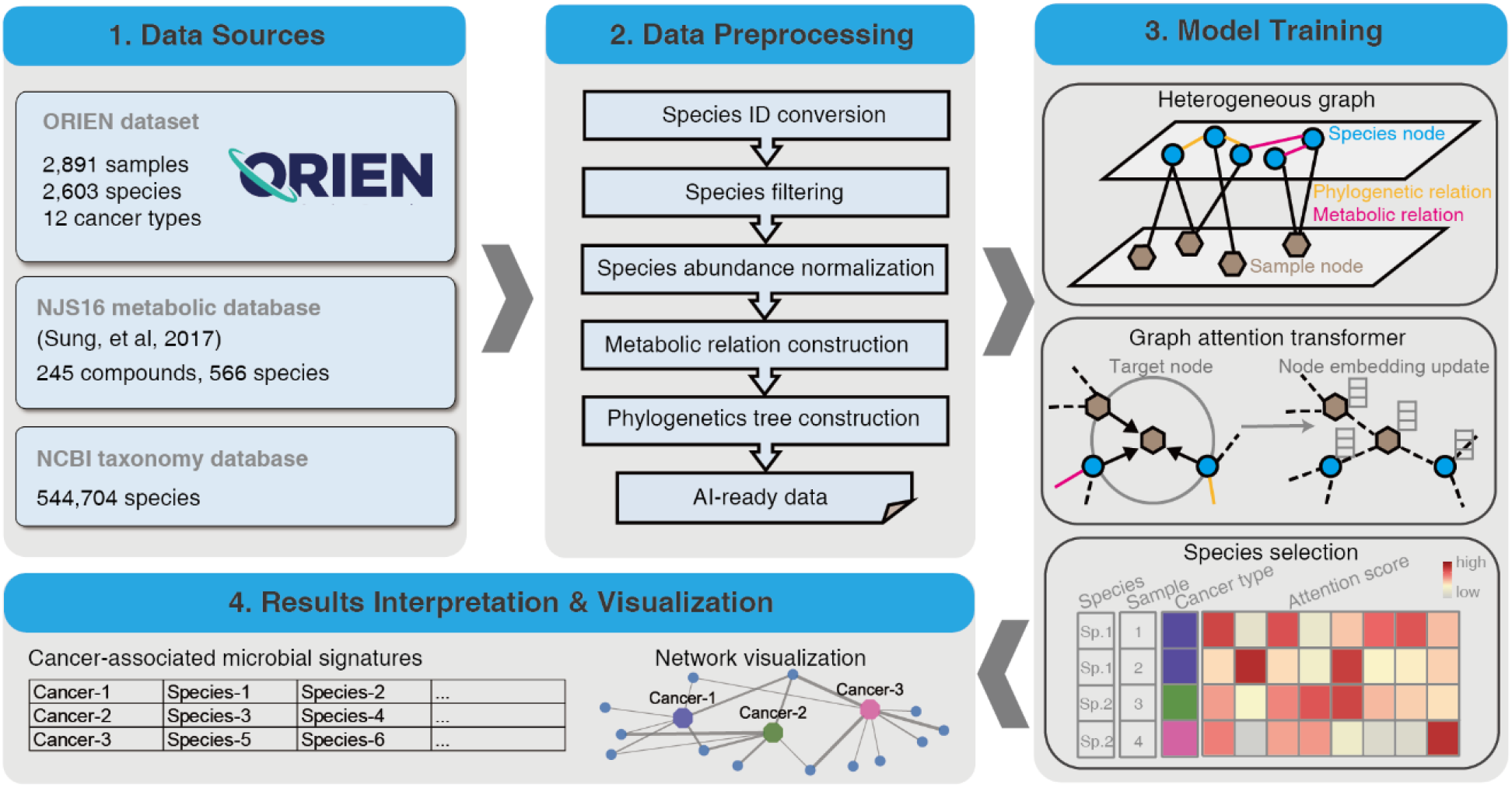
Overview of the MEGA workflow. Four main steps were included in carrying out model training and biological gene network inference. MEGA uses ORIEN datasets and two database dependencies as the data sources. Preprocessing steps are employed to generate artificial intelligence (AI)-ready data for graph neural network training. After deep learning model training, the cancer-associated microbial signatures were selected based on the attention scores of each species at the sample level. The final results of the identified cancer-associated microbial communities have been provided in a tabular format and are available for additional visualization.

### Data Preprocessing

We initially converted the organism’s name to a standard taxonomy ID using the *taxizedb* package. Species were filtered by removing those that expressed less than 0.1% of the total species. After filtering, 2,266 species were obtained. To normalize the microbiome matrix, we scaled the values in each sample of the matrix that summed to 1. This method ensures that the contribution of each feature to the total sum is proportional to its relative abundance in the sample. We used the normalized matrix as the basis for downstream analyses. Specifically, we generated the metabolic relationship network by comparing the total species list in the ORIEN matrix with the NJS16 metabolic database. In this network, an edge was placed between two species if they shared the same metabolic compound shown in the NJS16 database. Similarly, for the phylogenetic relation network, we compared the total species list in the ORIEN matrix with the NCBI taxonomy database, placing an edge that links two species if they share the same genus information. The processed data, including the normalized abundance matrix, metabolic relationship network, and phylogenetic relation network, served as artificial intelligence (AI)-ready data for model training **(see Figure 1 – Data Processing)**.

### Model Training

The main MEGA model was implemented in PyTorch (11) (v1.4.0) and was trained on an NVIDIA A100 graphics processing unit (GPU) for 50 epochs (approximately 15 minutes). We utilized our previously developed heterogeneous graph transformer model for model training (6). The input graph incorporates both species and sample nodes, along with the relations among them as edges. By capturing both neighbor and global topological features among samples and species, the model was able to construct sample-sample and species-species relations simultaneously. We used two autoencoders to generate the initial embeddings for the heterogeneous graph. This allowed the representation of each node as a dense vector, which can be used as input for the deep learning model. Meanwhile, we were able to reduce the dimensionality of each species and sample, resulting in an initial embedding size of 256 dimensions for all nodes in the graph. The complete heterogeneous graph embedding was subsequently passed to a graph attention transformer, which was trained to learn the relations between sample and species. MEGA adopts a heterogeneous multi-head attention mechanism to model the overall topological information (global relationships) and neighbor message passing (local relationships) on the heterogeneous graph. We used the Adam optimizer with a learning rate of 0.003 and default settings for other hyperparameters: n_hid=128, KL_COEF=0.00005, and THRES=3. The Focal Loss function was used to quantify the differences between the predicted cancer type labels and true cancer type labels. The learning rate was reduced by a factor of 0.5 when the evaluation metric stopped improving for 5 epochs. The heterogeneous graph representation learning facilitated the embedding of samples and species simultaneously using the transformer, yielding the attention score as an important training outcome. This score represents the importance of a source node to a target node. We extracted the attention scores from source nodes spanning from species to sample. A high attention score between a given species and a sample indicates that the species was highly represented in the sample. We leveraged this information to identify microbial signatures associated with specific cancer types. We accomplished this by counting the number of samples within the cancer type for each species with high attention scores. Species with a *p*-value less than 0.05 were considered to be significantly associated with the cancer type. These reliable microbial signatures were selected and served as the final output of MEGA **(see Figure 1 – Model Training)**.

### Results Interpretation and Visualization

The final output of MEGA is a tab-delimited list, where each row represents each cancer type followed by identified microbial signatures. The results can be visualized in UpSet plots (12) and Cytoscape networks (13). UpSet plots are a powerful visualization technique designed to display complex set data with more than 3 intersecting sets. This method provides an intuitive and comprehensive means of exploring the relationships between sets and their overlaps, allowing for a more nuanced interpretation of the underlying data. Cytoscape is a widely used open-source software platform that offers a suite of tools for the visualization, analysis, and modeling of complex networks. To leverage the strengths of Cytoscape’s capabilities, the RCy3 R package (14) was utilized to implement the network visualization aspect of MEGA. Through the use of Rcy3’s REST application programming interface (API), users can seamlessly access the full feature set of Cytoscape within the R programming environment. Users can import network works directly to Cytoscape with the predefined layout and theme using MEGA output files. The network comprises cancer-species nodes, with the thickness of the edges reflecting the attention weight scores. In addition, phylogenetic or metabolic relationships between species are represented by additional edges. This approach allows for a comprehensive and nuanced exploration of the relationships between cancer and species, providing valuable insights into the underlying biological processes and pathways involved. The attention weight scores, represented by the edge thickness, highlight the key connections and interactions within the network, enabling researchers to effectively identify potential targets for further study **(see Figure 1 – Results Interpretation & Visualization)**. Additional tutorials on generating both UpSet plots and Cytoscape networks can be found in the MEGA GitHub repository https://github.com/OSU-BMBL/MEGA.

### Implementation

MEGA was developed using Python 3.7.12 with PyTorch v1.4.0 and torch-geometric v1.4.3. The MEGA GPU mode was tested in CUDA v11.6 on a Red Hat Enterprise 7 Linux system 8.3, which featured 128-core AMD Epic central processing units (CPUs), NVIDIA A100-PCIE-80GB GPUs, and 1TB RAM. Similarly, the MEGA CPU mode was tested on the Ohio Supercomputer Center Pitzer cluster, which incorporated Intel Xeon Gold 6148 CPUs and 64GB RAM. MEGA was versioned and uploaded to the Python Package Index (PyPI) using Python-Versioneer, a tool that simplifies the management of version numbers in a software project. By subjecting the software to extensive testing in both GPU and CPU modes, we were able to ensure that MEGA functions effectively and efficiently across a range of computational architectures, ultimately providing users with a reliable and versatile tool.

## RESULTS

### MEGA Identifies Intratumoral Microbes from 12 Cancer Types in the ORIEN Dataset

Overall, MEGA is a deep learning package for identifying cancer-associated intratumoral microbes. It consists of 4 main steps: (1) Collect the ORIEN dataset, Human NJS16 metabolic database [2], and NCBI taxonomy database; (2) Preprocess ORIEN dataset as input for the deep learning model; (3) Train the graph attention transformer using a heterogeneous graph; and (4) Interpret cancer-associated intratumoral microbes. MEGA identified microbial communities that consist of 73 unique species from 12 cancer types in the ORIEN data **(see Figure 2 and Supplementary Table S1)**. Our analysis revealed that 15 species were shared across all 12 cancer types. Notably, 8 species were uniquely shared among COAD, rectum adenocarcinoma (READ), and other colorectal cancer (OtherCR). This group of 8 species represented the second-highest number of shared species across all intersections, and their shared presence is consistent with the fact that these cancers all originate in the large intestine, as in the case of colorectal cancer (CRC) **(see Supplementary Figure S1)**.

**Figure 2.**
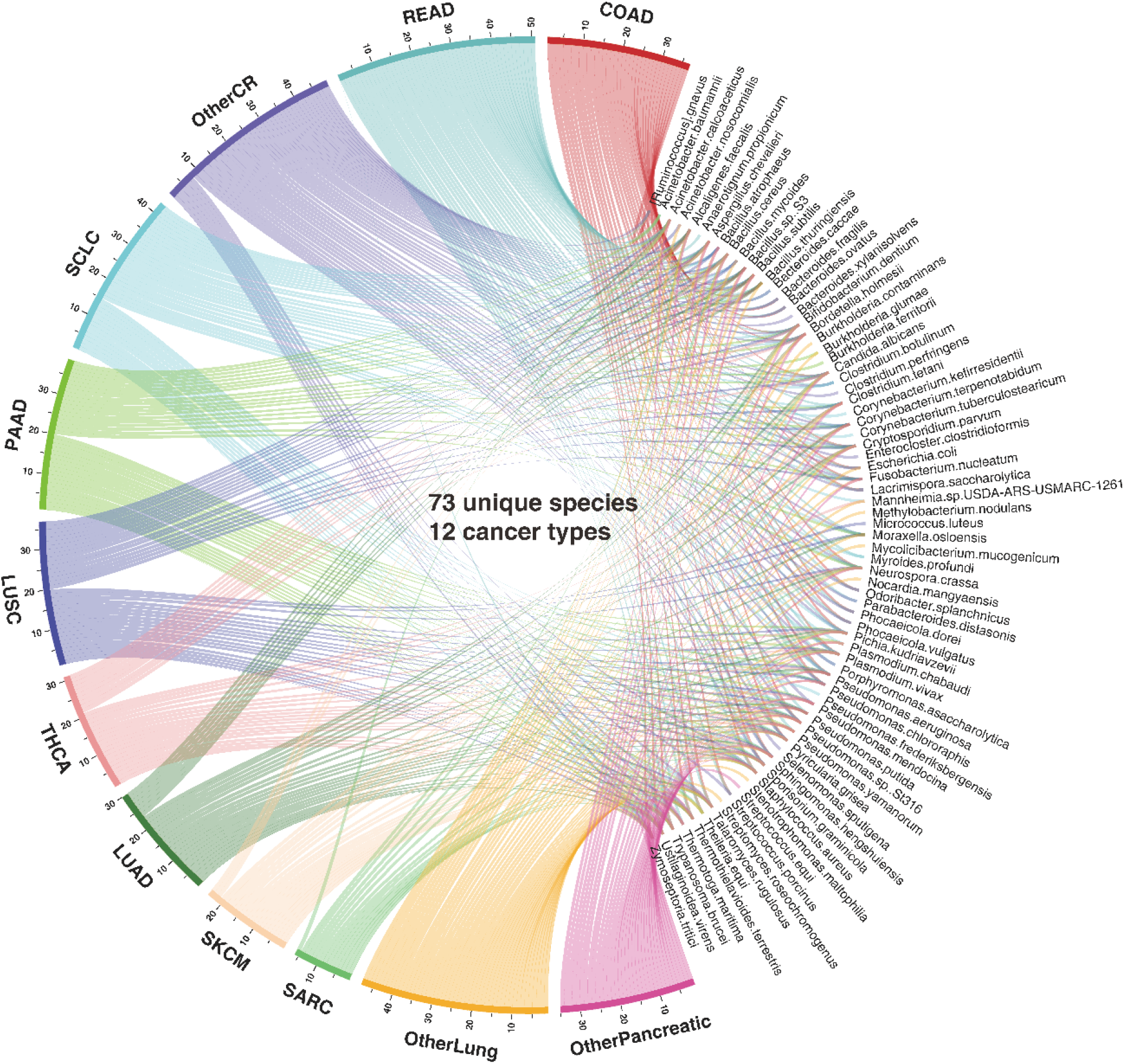
Circos plot representation of the distribution of identified species and cancer types. The segment length for each cancer type is proportional to the ratio of the total number of detected species within that cancer type, and individual ribbons are linked to their respective species. The cancer types are abbreviated as COAD (Colon Adenocarcinoma); LUAD (Lung Adenocarcinoma); LUSC (Lung Squamous Cell Carcinoma); OtherCR (Other colorectal cancer types not specified); OtherLung (Other lung cancer types not specified); OtherPancreatic (Other pancreatic cancer types not specified); PAAD (Pancreatic Adenocarcinoma); READ (Rectum Adenocarcinoma); SARC (Sarcoma); SCLC (Small Cell Lung Cancer); SKCM (Skin Cutaneous Melanoma); and THCA (Thyroid Carcinoma).

### MEGA Identifies Cancer-associated Microbes in Colon Adenocarcinoma and Thyroid Carcinoma

To demonstrate the data analysis and interpretation capabilities of MEGA, we focused on case studies in COAD and THCA. These cancers were chosen for their contrasting levels of attention within the tumor microbiome research community. COAD has been relatively well studied in relation to its associations with tumor microbes, whereas THCA has not yet received significant attention. By using these well-known cases as a benchmark, we validated the effectiveness and credibility of MEGA. COAD is a common malignant tumor in the digestive tract (15). Increased evidence suggests that intestinal microbiota was crucial in developing CRC (16). Our analysis revealed that 8 microbial species were uniquely shared among the CRC types COAD, READ, and OtherCR. These species are *Bacteroides fragilis, Ruminococcus gnavus, Bacteroides ovatus, Lacrimispora saccharolytica, Odoribacter splanchnicus, Phocaeicola dorei, Phocaeicola vulgatus*, and *Streptococcus porcinus*. Notably, 3 of these species—*Bacteroides fragilis, Ruminococcus gnavus*, and *Bacteroides ovatus*—were found to be consistent with previously validated experimental results (17-22). MEGA successfully identified these species by integrating metabolic and phylogenetic relationships in the model training process.

By integrating metabolic relationships, MEGA was able to capture the association even when a relatively low attention score is presented. For instance, *Fusobacterium nucleatum* shows high attention scores among the identified species in COAD, and its infection promotes CRC progression by changing the mucosal microbiota and colon transcriptome in a mouse model (17). *Ruminococcus gnavus* has a low attention score and the abundance was shown to have a significant negative correlation with CRC tumor numbers and disease score (18). *Fusobacterium nucleatum* and *Ruminococcus gnavus* shared the same compound C00270: N-Acetylneuraminate acid, where the intercellular adhesive events may play an important role in tumor angiogenesis, metastasis, and growth control in COAD (19). *Ruminococcus gnavus* also shared the same compound C01019: L-Fucose with *Bacteroides fragilis*. Recent studies found that *Bacteroides fragilis* toxin can contribute to COAD formation (20), while fucose-bound liposomes carrying anticancer drugs could serve as a new strategy for the treatment of CRC patients (21) **(see Figure 3A)**. THCA has increased substantially in many countries during the past few decades (23). The species related to compound C00422: Triacylglycerol, including *Pseudomonas aeruginosa* and *Staphylococcus aureus* were found in THCA groups. Recent studies suggest that elevated triglyceride levels may be a potential biomarker for identifying individuals at a higher risk of developing thyroid cancer (24) **(see Figure 3B)**. The full metabolic relationships for all 12 cancer types can be found in **Supplementary Table S2**.

**Figure 3.**
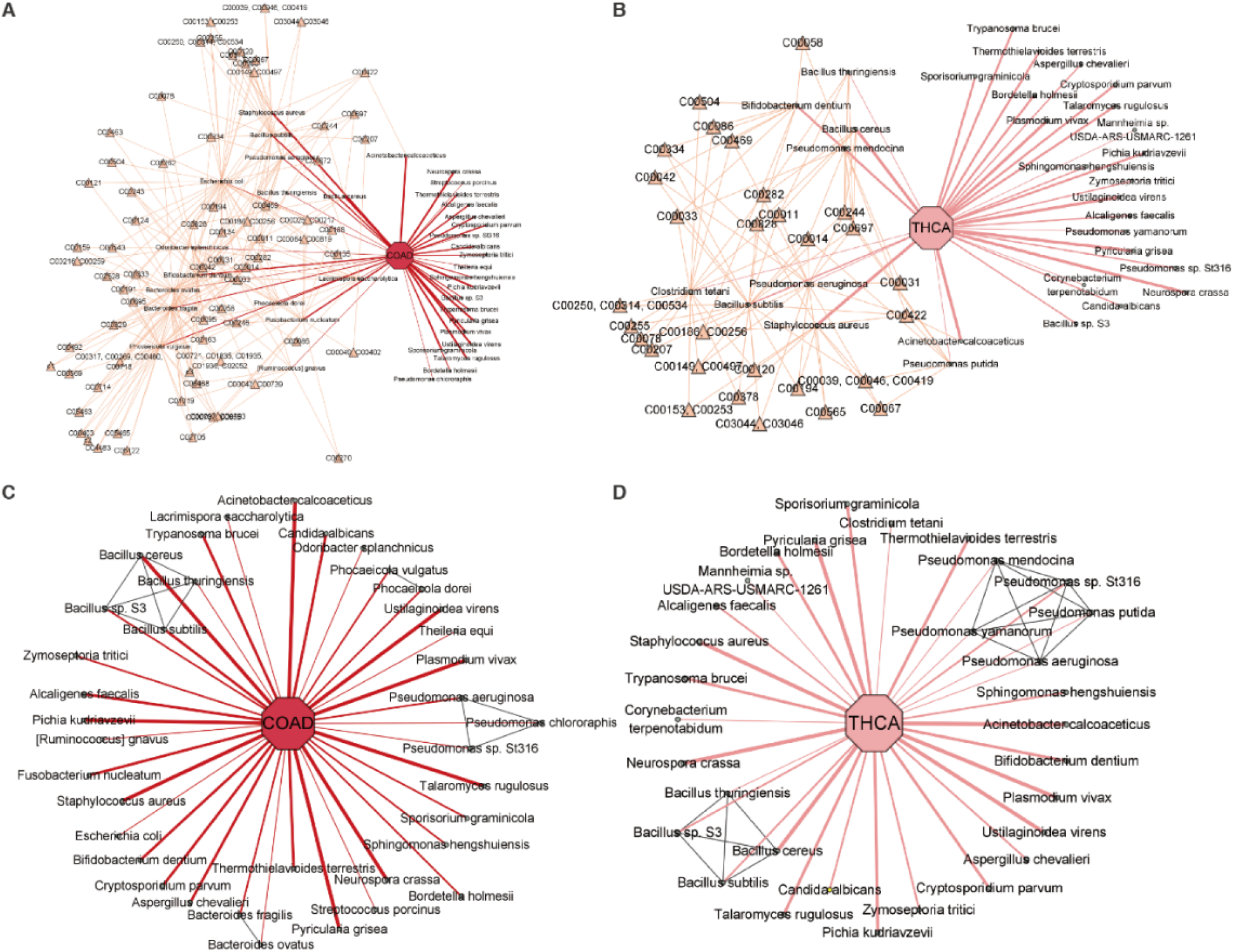
Network visualization of identified microbial communities in COAD and THCA. The cancer-type nodes were highlighted by an octagon shape, while the microbial species nodes were highlighted in a circle shape. The thickness of the edges in the network reflects the attention weight scores, indicating the strength of the relationship between the species and cancer. In addition, the metabolic compound nodes were highlighted with a yellow triangle shape, while the phylogenetic relationship edges were highlighted in gray. (**A)** COAD-associated microbes highlighted with metabolic compound. (**B)** THCA-associated microbes highlighted with metabolic compound. (**C)** COAD-associated microbes highlighted with phylogenetic relationships. **(D)** THCA-associated microbes highlighted with phylogenetic relationships.

By integrating phylogenetic relationships, MEGA was able to capture associations with relatively low attention scores. A previous study found that *Bacteroides ovatus* may be one of the dominant species in colon cancer (22). Although *Bacteroides ovatus* had a relatively low attention score, MEGA can identify it using the phylogenetic association with *Bacteroides fragilis*, which has a high attention score **(see Figure 3C)**. We found that *Pseudomonas mendocina, Pseudomonas putida*, and *Pseudomonas yamanorum* were uniquely identified in the *Pseudomonas* genus in THCA, in contrast to COAD. This aligns with the study showing the predominance of *Pseudomonas* in THCA **(see Figure 3D)** (25). The phylogenetic relationships for all 12 cancer types can be found in **Supplementary Table S3**.

## DISCUSSION

The development of MEGA represents a significant step forward in identifying and interpreting cancer-associated intratumoral microbes. The deep learning package presented in this study utilizes RNA-seq data from the ORIEN dataset to identify microbial signatures associated with 12 different cancer types. By leveraging the power of graph attention transformers, MEGA can capture both local and global topological features of the heterogeneous graph, resulting in a more comprehensive and nuanced understanding of the underlying biological processes and pathways involved. The application of MEGA to the ORIEN dataset has provided valuable insights into the role of intratumoral microbes in cancer. The analysis revealed 73 unique species associated with the 12 cancer types studied. Notably, 15 species were shared across all 12 cancer types, highlighting the potential importance of these microbes in cancer development and progression.

As a next step, we will further compare the cancer-associated intratumoral microbes identified from TCMA and ORIEN data using MEGA to provide a more comprehensive understanding of the role of intratumoral microbes in relation to cancer biology and host immunology. In the long run, the genotype-tissue expression (GTEx) data can be involved as control samples to identify relationships specific to tumors. In addition, applying MEGA to single-cell RNA-seq data could provide a more detailed understanding of the interactions between microbial communities and tumor cells at the cellular level. It may give us a new angle to characterize tumor heterogeneity based on intratumoral microbiome diversities. In conclusion, the development of MEGA represents an important advance in identifying cancer-associated intratumoral microbes. Our analysis of ORIEN data using MEGA revealed the presence of unique microbial signatures in specific cancer types, which may provide new targets for therapeutic intervention.

## SUPPLEMENTARY DATA

Supplementary Table S1. Identified microbial signatures with normalized attention weights.

Supplementary Table S2. Metabolic compounds relationships microbial signatures.

Supplementary Table S3. Phylogenetic relationships of microbial signatures.

## ACKNOWLEDGMENTS

This work was supported by the Pelotonia Institute of Immuno-Oncology (PIIO). The content is solely the responsibility of the authors and does not necessarily represent the official views of the PIIO. The authors acknowledge the support and resources of the Ohio Supercomputer Center (PAS1695, PCON0005). We would like to thank Angela Dahlberg, Editor, Division of Medical Oncology at The Ohio State University Comprehensive Cancer Center, for editing and proofreading the manuscript.

## REFERENCES

1. Cho, I. and Blaser, M.J. (2012) The human microbiome: at the interface of health and disease. Nat Rev Genet, 13, 260–270.

2. Chen, Y., Wu, F.H., Wu, P.Q., Xing, H.Y. and Ma, T. (2022) The Role of The Tumor Microbiome in Tumor Development and Its Treatment. Front Immunol, 13, 935846.

3. Hoyd, R., Wheeler, C.E., Liu, Y., Singh, M.J., Muniak, M., Denko, N., Carbone, D., Mo, X. and Spakowicz, D. (2022) Exogenous sequences in tumors and immune cells (exotic): a tool for estimating the microbe abundances in tumor RNAseq data.

4. Hu, Z., Dong, Y., Wang, K. and Sun, Y. (2020) Heterogeneous Graph Transformer.

5. Ma, A., Wang, X., Li, J., Wang, C., Xiao, T., Liu, Y., Cheng, H., Wang, J., Li, Y., Chang, Y. et al.. (2023) Single-cell biological network inference using a heterogeneous graph transformer. Nat Commun, 14, 964.

6. Liu, Z., Sun, Y., Ma, A., Wang, X., Xu, D., Spakowicz, D., Ma, Q. and Liu, B. (2023) An explainable graph neural framework to identify cancer-associated intratumoral microbial communities. bioRxiv, 2023.2004.2016.537088.

7. Dohlman, A.B., Arguijo Mendoza, D., Ding, S., Gao, M., Dressman, H., Iliev, I.D., Lipkin, S.M. and Shen, X. (2021) The cancer microbiome atlas: a pan-cancer comparative analysis to distinguish tissue-resident microbiota from contaminants. Cell Host Microbe, 29, 281–298 e285.

8. Dalton, W.S., Sullivan, D., Ecsedy, J. and Caligiuri, M.A. (2018) Patient Enrichment for Precision-Based Cancer Clinical Trials: Using Prospective Cohort Surveillance as an Approach to Improve Clinical Trials. Clin Pharmacol Ther, 104, 23–26.

9. Sung, J., Kim, S., Cabatbat, J.J.T., Jang, S., Jin, Y.-S., Jung, G.Y., Chia, N. and Kim, P.-J. (2017) Global metabolic interaction network of the human gut microbiota for context-specific community-scale analysis. Nature Communications, 8, 15393.

10. Schoch, C.L., Ciufo, S., Domrachev, M., Hotton, C.L., Kannan, S., Khovanskaya, R., Leipe, D., Mcveigh, R., O’Neill, K., Robbertse, B. et al.. (2020) NCBI Taxonomy: a comprehensive update on curation, resources and tools. Database, 2020, baaa062.

11. Paszke, A., Gross, S., Massa, F., Lerer, A., Bradbury, J., Chanan, G., Killeen, T., Lin, Z., Gimelshein, N., Antiga, L. et al.. (2019) PyTorch: An Imperative Style, High-Performance Deep Learning Library.

12. Lex, A., Gehlenborg, N., Strobelt, H., Vuillemot, R. and Pfister, H. (2014) UpSet: Visualization of Intersecting Sets. IEEE transactions on visualization and computer graphics, 20, 1983–1992.

13. Shannon, P., Markiel, A., Ozier, O., Baliga, N.S., Wang, J.T., Ramage, D., Amin, N., Schwikowski, B. and Ideker, T. (2003) Cytoscape: A Software Environment for Integrated Models of Biomolecular Interaction Networks. Genome Research, 13, 2498–2504.

14. Gustavsen, J.A., Pai, S., Isserlin, R., Demchak, B. and Pico, A.R. (2019). F1000Research.

15. Xie, Y.-H., Chen, Y.-X. and Fang, J.-Y. (2020) Comprehensive review of targeted therapy for colorectal cancer. Signal Transduction and Targeted Therapy, 5, 1–30.

16. Lucas, C., Barnich, N. and Nguyen, H.T.T. (2017) Microbiota, Inflammation and Colorectal Cancer. International Journal of Molecular Sciences, 18, 1310.

17. Wu, N., Feng, Y.-Q., Lyu, N., Wang, D., Yu, W.-D. and Hu, Y.-F. (2022) Fusobacterium nucleatum promotes colon cancer progression by changing the mucosal microbiota and colon transcriptome in a mouse model. World Journal of Gastroenterology, 28, 1981–1995.

18. Alrafas, H.R., Busbee, P.B., Chitrala, K.N., Nagarkatti, M. and Nagarkatti, P. (2020) Alterations in the Gut Microbiome and Suppression of Histone Deacetylases by Resveratrol Are Associated with Attenuation of Colonic Inflammation and Protection Against Colorectal Cancer. Journal of Clinical Medicine, 9, 1796.

19. Dimitroff, C.J., Pera, P., Dall’Olio, F., Matta, K.L., Chandrasekaran, E.V., Lau, J.T. and Bernacki, R.J. (1999) Cell surface n-acetylneuraminic acid alpha2,3-galactoside-dependent intercellular adhesion of human colon cancer cells. Biochemical and Biophysical Research Communications, 256, 631–636.

20. Cheng, W.T., Kantilal, H.K. and Davamani, F. (2020) The Mechanism of Bacteroides fragilis Toxin Contributes to Colon Cancer Formation. The Malaysian Journal of Medical Sciences : MJMS, 27, 9–21.

21. Osuga, T., Takimoto, R., Ono, M., Hirakawa, M., Yoshida, M., Okagawa, Y., Uemura, N., Arihara, Y., Sato, Y., Tamura, F. et al.. (2016) Relationship Between Increased Fucosylation and Metastatic Potential in Colorectal Cancer. JNCI: Journal of the National Cancer Institute, 108, djw210.

22. He, T., Cheng, X. and Xing, C. (2021) The gut microbial diversity of colon cancer patients and the clinical significance. Bioengineered, 12, 7046–7060.

23. Kitahara, C.M. and Sosa, J.A. (2016) The changing incidence of thyroid cancer. Nature Reviews Endocrinology, 12, 646–653.

24. Alkurt, E.G., Şahin, F., Tutan, B., Canal, K. and Turhan, V.B. (2022) The relationship between papillary thyroid cancer and triglyceride/glucose index, which is an indicator of insulin resistance. European Review for Medical and Pharmacological Sciences, 26, 6114–6120.

25. Yuan, L., Yang, P., Wei, G., Hu, X.e., Chen, S., Lu, J., Yang, L., He, X. and Bao, G. (2022) Tumor microbiome diversity influences papillary thyroid cancer invasion. Communications Biology, 5, 1–9.

