## Supplementary Information for "A bioinformatics tool for identifying intratumoral microbes from the ORIEN dataset"

Software source code: https://github.com/OSU-BMBL/MEGA

Supplementary Figure S1: Upset plot of overall identified species in 12 cancer types.

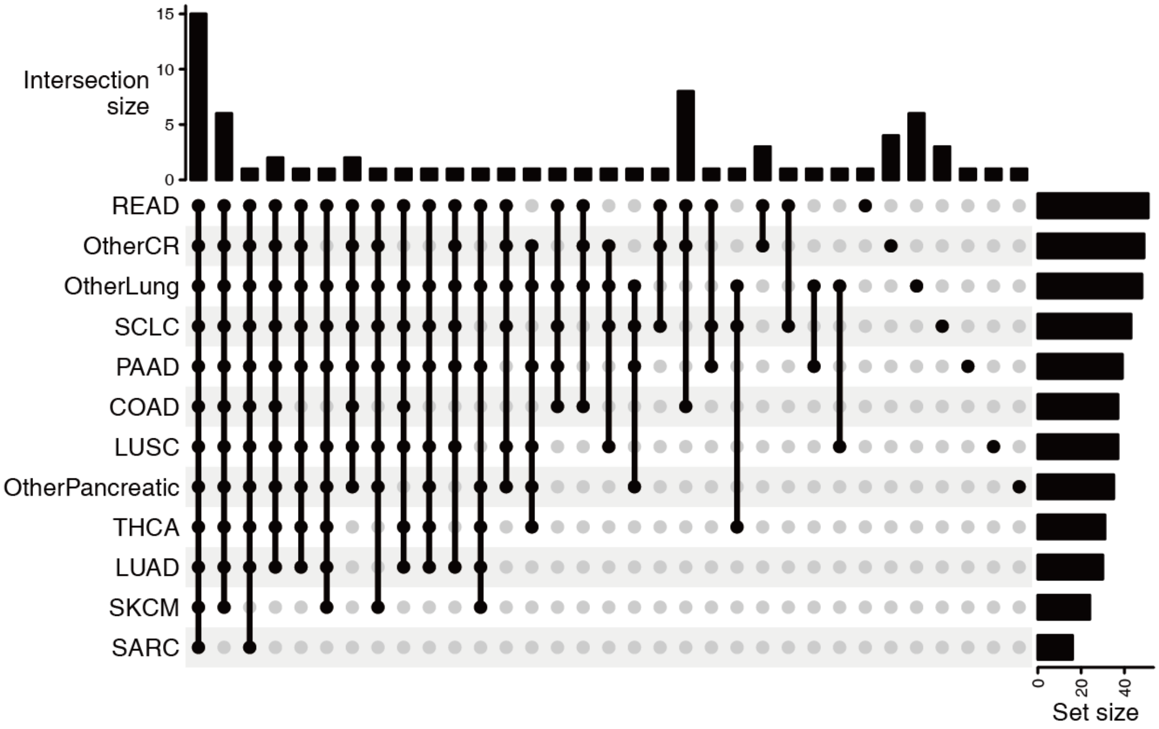


**Supplementary Figure S1**: Upset plot showing the overlapping relationships of the identified species among each cancer type. The rows represent the species sets within each cancer types, while the columns represent the intersections between these sets. The bar charts on the sides depict the size of the sets. The cancer types are abbreviated as: COAD (Colon Adenocarcinoma), LUAD (Lung Adenocarcinoma), LUSC (Lung Squamous Cell Carcinoma), OTHERCR (Other colorectal cancer types not specified), OtherLung (Other lung cancer types not specified), OtherPancreatic (Other pancreatic cancer types not specified), PAAD (Pancreatic Adenocarcinoma), READ (Rectum Adenocarcinoma), SARC (Sarcoma), SCLC (Small Cell Lung Cancer), SKCM (Skin Cutaneous Melanoma), and THCA (Thyroid Carcinoma).
